# Amino acids whose intracellular levels change most during aging alter chronological lifespan of fission yeast

**DOI:** 10.1101/2020.08.18.256164

**Authors:** Charalampos Rallis, Michael Mülleder, Graeme Smith, Yan Zi Au, Markus Ralser, Jürg Bähler

## Abstract

Amino acid deprivation or supplementation can affect cellular and organismal lifespan, but we know little about the role of concentration changes in free, intracellular amino acids during aging. Here, we determine free amino-acid levels during chronological aging of non-dividing fission yeast cells. We compare wild-type with long-lived mutant cells that lack the Pka1 protein of the protein kinase A signalling pathway. In wild-type cells, total amino-acid levels decrease during aging, but much less so in *pka1* mutants. Two amino acids strongly change as a function of age: glutamine decreases, especially in wild-type cells, while aspartate increases, especially in *pka1* mutants. Supplementation of glutamine is sufficient to extend the chronological lifespan of wild-type but not of *pka1Δ* cells. Supplementation of aspartate, on the other hand, shortens the lifespan of *pka1Δ* but not of wild-type cells. Our results raise the possibility that certain amino acids are biomarkers of aging, and their concentrations during aging can promote or limit cellular lifespan.

## Introduction

Dietary restriction extends lifespan and decreases age-related pathologies (1,2). These benefits may reflect protein or amino-acid restrictions rather than overall calorie intake (3,4). Restriction or supplementation of certain amino-acids affect lifespan from yeast to mouse (5). In budding yeast, isoleucine, threonine and valine extend the chronological lifespan (CLS, the time post-mitotic cells remain viable in stationary phase) (6). In other yeast strains, serine, threonine and valine decrease the CLS (7), while removal of asparagine extends the CLS (8). In mice, lifespan is extended by the branched chain amino acids (BCAAs: leucine, isoleucine and valine) (9), while in flies, limitation of BCAAs extends lifespan in a dietary-nitrogen-dependent manner (10). In mice, a tryptophan-restricted diet extends lifespan (11), while a methionine-restricted diet extends lifespan and delays age-related phenotypes (12). In flies, methionine restriction extends lifespan (13). Similarly, in budding yeast, methionine restriction prolongs the CLS whilst methionine addition shortens it (14). A comprehensive analysis of amino-acid effects on lifespan has been undertaken in worms, where lifespan extends upon individual supplementation of 18 amino acids (15). These results indicate that amino acids can exert both pro- and anti-aging effects. In budding yeast, intracellular amino-acid content gradually decreases during chronological aging (16). What is missing, however, is a global quantitative analysis of intracellular amino acids in normal and long-lived cells during cellular aging.

In fission yeast (*Schizosaccharomyces pombe*), the protein kinase A (PKA) glucose-sensing pathway promotes chronological aging (19). Here we perform quantitative amino-acid profiling during aging in wild-type and long-lived *pka1* mutant cells of *S. pombe*, which are deleted for the catalytic subunit of PKA. We show that intracellular amino-acid pools progressively decrease with age. This effect is less pronounced in long-lived cells. Glutamine is depleted faster than the other amino acids, while aspartate increases during aging. Notably, supplementation of these two amino acids is sufficient to alter lifespan in wild-type or *pka1* mutant cells. Our results suggest both a correlative and causal relationship between longevity and intracellular amino acids.

## Results

### Quantitative amino-acid analysis during cellular aging

We used liquid chromatography selective reaction monitoring (19) to quantify the free, intracellular pools for 19 of the 20 canonical proteogenic amino acids (cysteine was excluded as it is readily oxidized) during chronological aging of *S. pombe* wild-type and long-lived *pka1Δ* deletion-mutant cells. Cells were grown in minimal medium, because intracellular amino acids accumulate in rich medium (20). We collected cells from three independent biological experiments at three timepoints each: at Day 0, when cells enter stationary phase and both strains show 100% cell viability; at Day 5, when wild-type and *pka1Δ* strains show 50% and 87% viability, respectively; and at Day 8, when wild-type and *pka1Δ* strains show 20% and 50% viability, respectively (Figure 1A).

**Figure 1.**
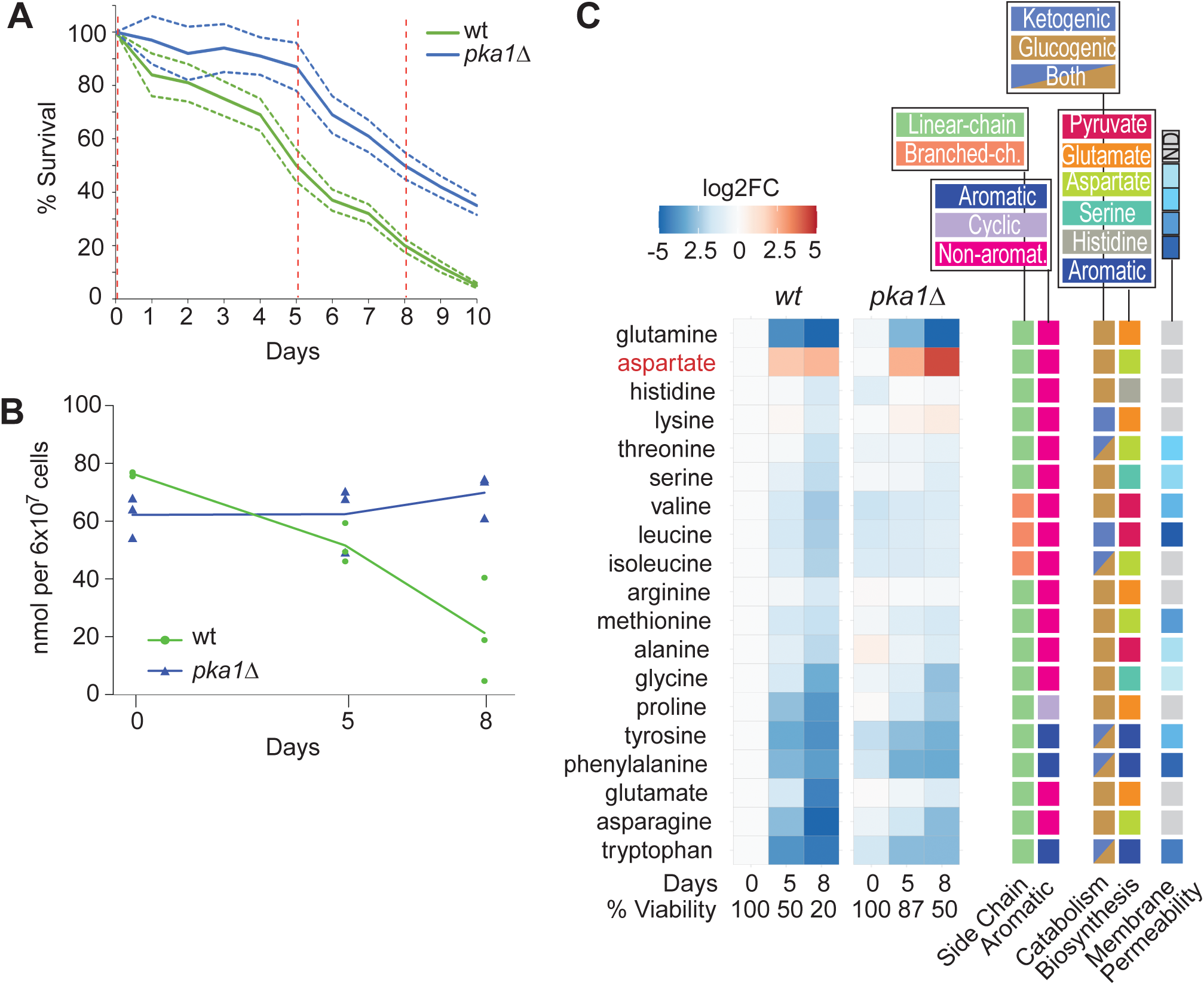
Quantitative measurements of free, intracellular amino acids in chronologically aging cells. **A**. CLS assays for wild-type and *pka1Δ* cells, performed in triplicate with each biological replicate measured in three technical repeats. Average lifespans are shown (solid lines) together with standard deviations (dotted lines). Hatched red lines indicate timepoints for amino-acid profiling. **B**. Total amino-acid concentrations in wild-type and *pka1Δ* cells at different aging timepoints as indicated. **C**. Heatmap showing quantitation of individual amino acids. The log2 fold-change values relative to 100% viability of wild-type cells (Day 0) were clustered using the R package pheatmap with standard settings. Averages of three timepoints with three independent biological repeats each are shown.

We determined absolute concentrations of intracellular amino acids in nmol/6×10^7^ cells (eTable 1). The amino-acid quantities, obtained from non-dividing cells, differed from those previously reported for fission yeast which have been obtained from rapidly proliferating cells (21,22). At the onset of stationary phase, the total amino-acid concentrations were lower in *pka1Δ* than in wild-type cells (Figure 1B). Accordingly, the individual amino-acid concentrations were lower or similar in *pka1Δ* than in wild-type cells at that stage (Figure 1C; Figure 2). The concentration differences were particularly pronounced for the BCAAs and aromatic amino acids (phenylalanine, tryptophan, tyrosine) (Figure 1C; Figure 2).

**Figure 2.**
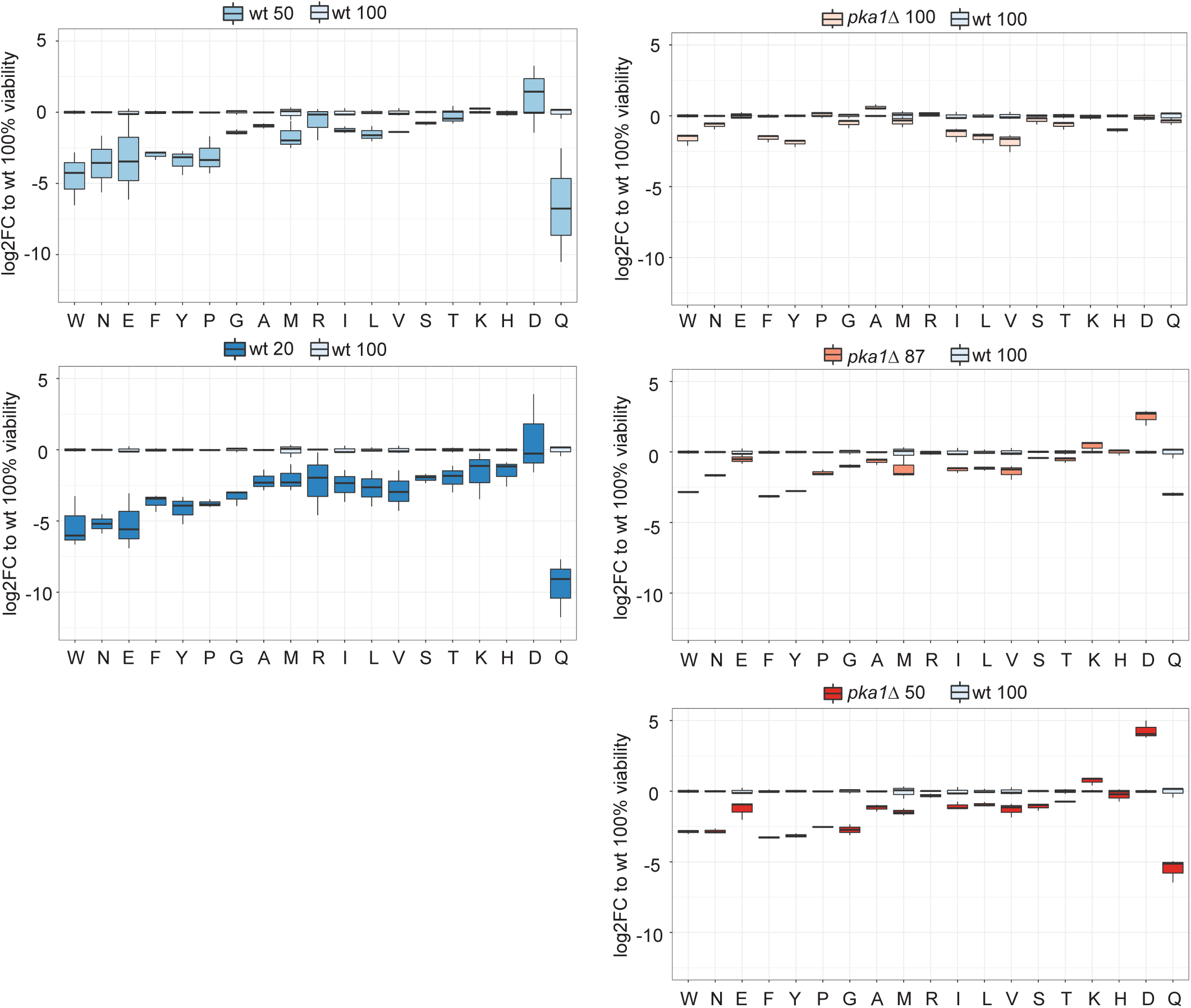
Changing amino-acid concentrations during aging. Normalised concentration values of 19 amino acids in wild-type cells (left) and *pka1Δ* cells (right) during aging (top to bottom). For each condition, three independent samples were analysed. Concentrations are presented relative to mean value for given amino acid in wild-type cells at 100% viability (wt 100). Boxplots were created with R boxplot and default settings. Numbers next to strain at top of each graph represent % cell viability.

During chronological aging, the concentration of total free amino acids declined in wild-type but less so in *pka1Δ* cells, even rising slightly at the last timepoint (Figure 1B). This rise primarily reflected an increase in the most abundant amino-acid, lysine, which compensated for the decline in other amino acids (Figure 2). In wild-type samples, the variation in free amino acids between experimental repeats noticeably increased during aging, in contrast to *pka1Δ* samples (Figure 1B; Figure 2). This result raises the possibility that increased variation of amino-acid concentrations, reflecting less tight metabolic regulation, is a feature of wild-type aging cultures.

Wild-type cells showed a general decrease in amino acids during aging, apart from aspartate (Figure 1C; Figure 2). The BCAAs were initially present at ∼3-fold higher levels in wild-type cells before dropping to about half the level of *pka1Δ* cells (Figure 1C; Figure 2). No correlations were evident between aging-dependent concentration changes of amino acids and their chemical/physical features or metabolic pathways (Figure 1C). This result suggests that changes in amino-acid concentrations are not driven by limitation of precursor molecules. Likewise, there was no clustering based on glucogenic amino acids, which can be converted into glucose through gluconeogenesis (Figure 1C). This result suggests that any need for gluconeogenesis under glucose depletion does not greatly affect free amino-acid composition. Reassuringly, there was also no clustering based on membrane permeability (Figure 1C), as this suggests that the results were not biased by amino acids leaking from non-viable cells during the aging timecourse.

Notably, the amino-acid profiles of wild-type and *pka1Δ* cells were similar overall at corresponding aging timepoints, when both strains showed 50% viability (Figure 1B,C; Figure 2). This result suggests that similar metabolic changes occur in aging wild-type and mutant cells, but that these changes are delayed in the long-lived mutant cells. Some amino acids, however, showed distinct patterns in wild-type and *pka1Δ* cells, including lysine, glutamate, glutamine and aspartate (Figure 1C; Figure 2). Glutamine and aspartate showed particularly striking profiles during aging. Glutamine rapidly and strongly decreased during aging, more pronounced in wild-type cells (Figure 1C; Figure 2). Aspartate, on the other hand, was the only amino acid that increased during aging in wild-type cells, and this increase was more pronounced in *pka1Δ* cells (Figure 1C; Figure 2). These results pointed to glutamine and aspartate as markers for aging in *S. pombe* and raised the possibility that these amino acids directly contribute to cellular lifespan.

### Glutamine supplementation extends lifespan of wild-type cells

To examine the effect of glutamine on the CLS of wild-type and *pka1Δ* cells, we grew cultures to stationary phase in EMM2 minimal medium. During chronological aging, we replaced the medium with either EMM2 (control) on Day 1 or with EMM2 plus glutamine on either Day 1 (viability 84±8% for wild-type and 97±9% for *pka1Δ*) or Day 5 (viability 50±6% for wild-type and 87±8% for *pka1Δ*). These manipulations did not affect cell numbers or total protein levels of the aging cultures (eFigure 1). In wild-type cells, glutamine supplementation significantly extended the CLS, both when added on Day 1 or 5 (Figure 3A,B). Glutamine supplementation at Day1 extended the medial CLS from 5 to 6.5 days. Although Day 5 was close to the median lifespan of wild-type cells, glutamine still extended the CLS even at this late stage (Figure 3A,B). These results indicate that glutamine is beneficial for viability of aging wild-type cells. In *pka1Δ* cells, on the other hand, glutamine supplementation at either Day 1 or 5 had no effect on lifespan, with the median CLS remaining ∼8 days (Figure 3C,D). We conclude that glutamine addition during cellular aging promotes longevity in wild-type cells, but not in the long-lived *pka1Δ* cells which maintain relatively higher glutamine levels during aging (Figure 2).

**Figure 3.**
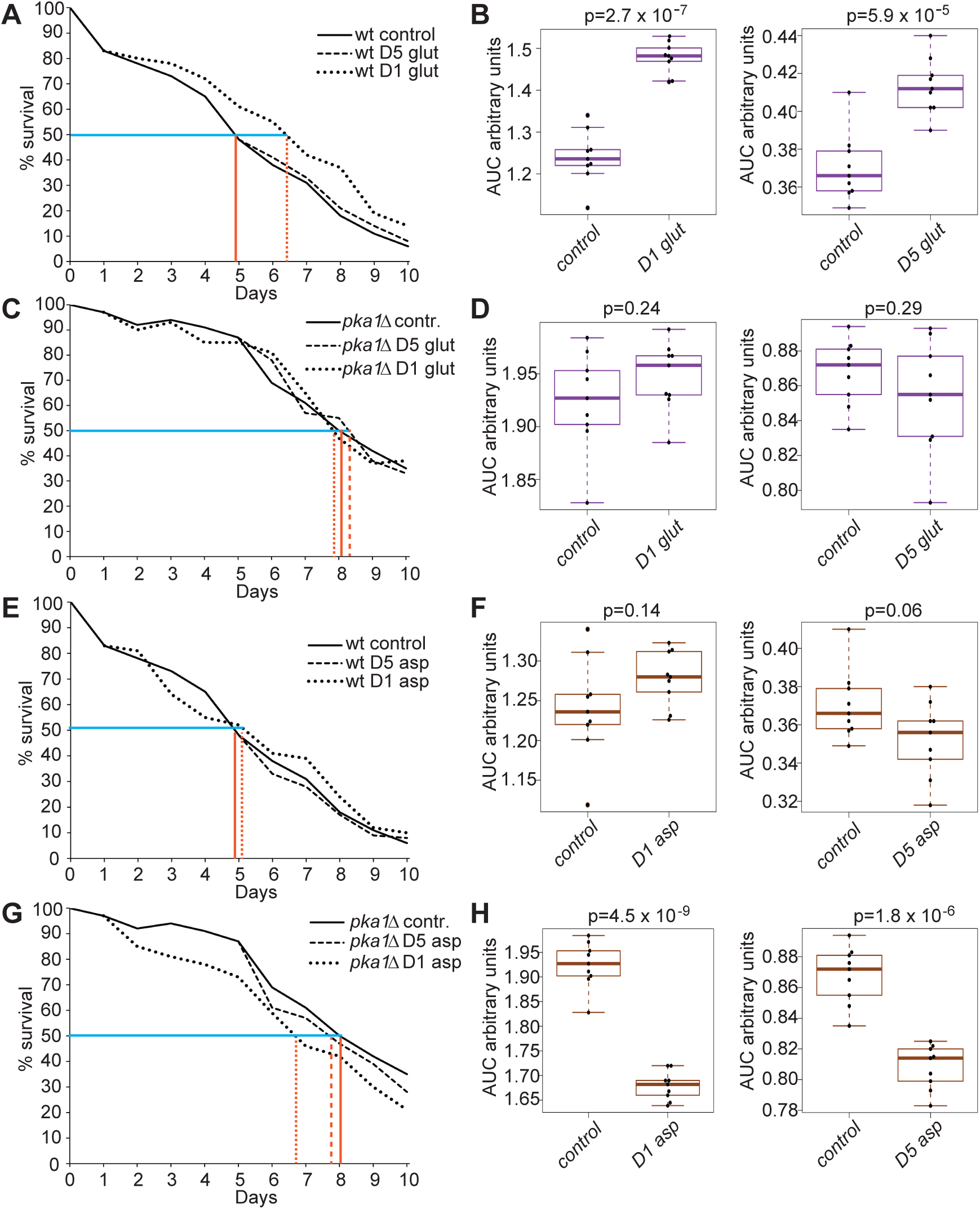
Effects of amino-acid supplementation on CLS. **A**. CLS assays (average of three biological repeats with three technical repeats each) for wild-type cells with and without glutamine supplementation at Days 1 and 5 as indicated. Median CLS in control cells (horizontal orange line) and treated cells (dotted orange line) are shown. **B**. Areas under curve (AUC) of CLS assays in A as indicated (left panel: AUCs from Day1, right panel: AUCs from Day 5). The p values (t-test) indicate significance of difference in CLS triggered by glutamine supplementation. **C**. CLS assays as in A for *pka1Δ* cells with and without glutamine supplementation at Days 1 and 5. **D**. AUC of CLS assays in C, as described in B. **E**. CLS assays as in A for wild-type cells with and without aspartate supplementation at Days 1 and 5. **F**. AUC of CLS assays in E, as described in B. **G**. CLS assays as in A for *pka1Δ* cells with and without aspartate supplementation at Days 1 and 5. **H**. AUC of CLS assays in G, as described in B.

### *Aspartate supplementation shortens lifespan of* pka1Δ *cells*

To examine the effect of aspartate on the CLS of wild-type and *pka1Δ* cells, we performed the same experiment as with glutamine, but supplementing aspartate to chronologically aging cultures. Again, these manipulations did not affect cell numbers or total protein levels of the aging cultures (eFigure 1). In wild-type cells, aspartate supplementation at either Day 1 or 5 had no effect on the CLS (Figure 3E,F). In *pka1Δ* cells, on the other hand, aspartate led to a significant shorter CLS, both when applied on Day 1 or 5 (median CLS shortened from 8 to 6.7 or 7.7 days, respectively); however, the median CLS remained longer than that of wild-type cells (Figure 3G,H). We conclude that aspartate addition during cellular aging has no effect in wild-type cells but shortens the CLS in *pka1Δ* cells where aspartate naturally strongly increases during aging (Figure 2).

## Discussion

We report intracellular amino-acid concentrations in *S. pombe* as a function of both chronological aging and genetic background. Our results show an overall decrease in amino acids during chronological aging, especially in wild-type cells. Such a decrease has also been observed in budding yeast (23). We cannot exclude the possibility that some dead cells contributed to these amino-acid measurements, although amino acids with high membrane permeability do not increase with age and thus do not preferentially leak from dead cells. The amount and composition of most amino acids reflect the ‘biological age’ of wild-type and long-lived *pka1Δ* cells, being similar in cells of similar viability rather than similar chronological age. Hence, 5 day old wild-type cells feature a similar amino-acid signature as 8-day old *pka1Δ* cells, corresponding to the times when both strains show 50% viability. Such amino-acid signatures might therefore serve as aging biomarkers for *S. pombe*.

Glutamine and aspartate show the most distinct profiles during aging, with a strong decrease of glutamine, especially in wild-type cells, and a strong increase of aspartate, especially in *pka1Δ* cells. The antagonistic changes in glutamine and aspartate are probably linked. During glucose deprivation, yeast cells turn to glutamine and glutamate for energy by making aspartate via glutaminolysis (24). Increased aspartate levels in aging *pka1Δ* cells could reflect that long-lived cells feature more efficient glutaminolysis, although alanine, another product of glutaminolysis, did not show the same trend. Induced glutaminolysis in long-lived cells could explain why aspartate did not increase lifespan in *pka1Δ* cells which naturally feature high aspartate levels.

Glutamine supplementation promotes longevity of wild-type but not of long-lived *pka1Δ* cells. Aspartate supplementation, on the other hand, shortens the lifespan of *pka1Δ* but not of wild-type cells. These amino acids also affect lifespan in worms (15): glutamine at high doses extends lifespan but at a lower dose shortens lifespan, while aspartate shortens lifespan. Glutamine and aspartate both affect mitochondrial functions which might mediate their lifespan effects. Glutamine, derived from the Krebs-cycle metabolite alpha-ketoglutarate, is one of the amino acids recently shown to become limited when blocking respiration in fermentatively growing *S. pombe* cells (21). Further experiments will provide mechanistic insights into the roles of glutamine and aspartate during aging. Nevertheless, our results suggest that decreased glutamine levels during aging not only correlate with aging in wild-type and *pka1Δ* cells but directly contribute to aging, which can be ‘cured’ by glutamine addition in wild-type cells. These findings highlight the metabolic complexity of aging and its relationship with nutrient-sensing pathways like PKA. Interestingly, decreased glutamine levels are associated with aging also in budding yeast, rats and humans (16,25), suggesting that conserved cellular processes are involved in this phenomenon.

## Methods

### Strains and media

Strain *972 h*^*-*^ was used as the reference wild-type. The *pka1* deletion mutant (*pka1::kanMX4 h-*) was generated with standard methods (26). Strains were cultured in EMM2 minimal medium for mass spectrometry sample acquisition, chronological aging and amino-acid supplementation experiments. Liquid cultures were grown at 32°C with shaking at 170rpm.

### Quantitative amino-acid profiling

Amino-acid analysis was performed using hydrophilic interaction chromatography-tandem mass spectrometry (HILIC-MS/MS) as described (22). Cell numbers were determined with a Beckman Z-series coulter counter to ensure equal cell amounts for extractions. The metabolites were extracted as described (22). Identification of 19 proteogenic amino acids was obtained by comparison of retention time and fragmentation patterns with commercial standards. Comparison against a standard curve produced from serial dilution of these standards allowed quantification of the free amino acids. The analysis was undertaken using an Agilent Infinity 1290 LC system with ACQUITY UPLC BEH amide columns (Waters Corporation, Manchester, UK) (pore diameter 130Å, particle size 1.7μm, internal diameter 2.1, column length 100mm) coupled to an Agilent 6400 Series Triple Quadrupole LC/MS mass spectrometer operating in selected reaction monitoring mode. Cysteine was excluded from analysis due to its low stability. The co-eluting isomers threonine and homoserine were de-convolved using the homoserine-specific transition (m/z of 120->44).

### Amino acid supplementations

Aging cells were incubated in media containing amino acid supplements at Days 1 and 5, by removing the supernatant after centrifugation (1000rpm, 3min) followed by resuspension in EMM2, EMM2 with 20mM glutamine, or EMM2 with 20mM aspartate up to their original culture volume. Amino acids were not directly added to medium because of solubility issues.

### Chronological lifespan assays

CLS assays were performed as described (17). Error bars represent standard deviations (of 9 measurements), calculated from three independent cell cultures, with each culture measured three times at each timepoint. Areas under the curve (AUCs) were measured for all experimental repeats with ImageJ (27). We compared AUCs for the portion of lifespan curve after amino-acids supplementation as before this time lifespan curves were identical.

### Total protein measurements

We collected 10ml liquid cultures during exponential growth (OD_600_=0.5-1.0), the start of stationary phase (Day 0), and after amino-acid supplementation (Day 5). Cells were lysed using a Fastprep®-24 (6.5m/s, 60sec) with addition of 300μl lysis buffer and 0.5 mm diameter glass beads. Protein concentrations were quantified by a Pierce™ BCA Protein Assay Kit following the manufacturer’s protocol, and absorbance values were obtained using the Magellan™ Data Analysis software. Proteins were separated in 4-12% Bis-Tris gel (NuPAGE®), and protein bands were detected using Ponceau S staining solution (Sigma, P7170).

## Acknowledgement

We thank Shajahan Anver, Clara Correia-Melo and Stephan Kamrad for critical reading and valuable comments on the manuscript.

