## Supplemental Fig 1 and Table 1 for "Amino acids whose intracellular levels change most during aging alter chronological lifespan of fission yeast"

#### Appendix

**eFigure 1: No effects of amino acids on aging cell numbers or protein levels.**

**eTable 1: Quantitation of 19 free amino acids (nmol/6 x 10exp7 cells) during chronological aging of wild-type and *pka1* mutant cells.**

### eFigure 1

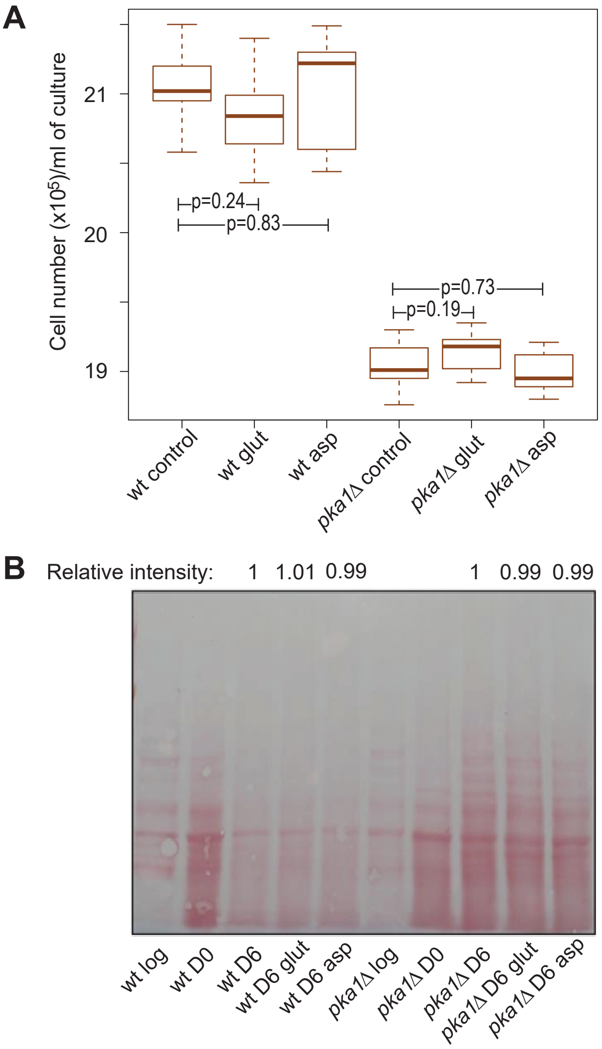


**e Figure1. No effects of amino acids on ageing cell numbers or protein levels.**
**A.** Cell numbers are unchanged within aged cultures following amino acid supplementation as indicated. Cell numbers were determined 24 hrs following change with media containing the supplemented amino acid.
**B.** Protein content is unchanged following amino acid supplementation as indicated.

**eTable 1: Quantitation of 19 free amino acids (in nmol/6 x 10exp7 cells) during chronological aging of wild-type and *pka1* mutant cells**

| Sample | ala | asp | glu | phe | gly | his | ile | lys | leu | met | asn | pro | gln | arg | ser | thr | val | trp | tyr |
|---|---|---|---|---|---|---|---|---|---|---|---|---|---|---|---|---|---|---|---|
| **wt_100_A** | 0.977 | 0.281 | 4.01 | 1.922 | 2.559 | 4.138 | 0.984 | 28.203 | 1.573 | 0.027 | 1.862 | 0.304 | 18.373 | 6.267 | 0.614 | 0.788 | 1.381 | 0.267 | 2.172 |
| **wt_100_B** | 0.994 | 0.289 | 3.956 | 1.958 | 2.537 | 4.619 | 1.003 | 27.09 | 1.637 | 0.018 | 2.07 | 0.307 | 18.779 | 6.078 | 0.664 | 0.866 | 1.499 | 0.291 | 2.342 |
| **wt_100_C** | 1.051 | 0.327 | 5.214 | 2.211 | 2.129 | 3.87 | 1.372 | 30.793 | 1.931 | 0.032 | 1.936 | 0.325 | 11.973 | 6.243 | 0.682 | 0.682 | 1.962 | 0.32 | 2.51 |
| **wt_50_A** | 0.585 | 2.875 | 4.251 | 0.294 | 1.039 | 3.927 | 0.564 | 32.32 | 0.881 | 0.017 | 0.623 | 0.096 | 2.833 | 7.224 | 0.398 | 0.452 | 0.603 | 0.041 | 0.355 |
| **wt_50_B** | 0.454 | 0.812 | 0.398 | 0.284 | 0.806 | 3.517 | 0.457 | 34.448 | 0.557 | 0.006 | 0.165 | 0.03 | 0.15 | 5.544 | 0.346 | 0.569 | 0.616 | 0.015 | 0.261 |
| **wt_50_C** | 0.526 | 0.11 | 0.062 | 0.198 | 0.881 | 3.969 | 0.403 | 35.722 | 0.414 | 0.004 | 0.04 | 0.016 | 0.011 | 1.585 | 0.393 | 1.058 | 0.621 | 0.003 | 0.11 |
| **wt_20_A** | 0.384 | 4.499 | 0.527 | 0.216 | 0.302 | 2.284 | 0.416 | 24.141 | 0.629 | 0.013 | 0.085 | 0.028 | 0.079 | 5.454 | 0.202 | 0.358 | 0.597 | 0.031 | 0.237 |
| **wt_20_B** | 0.203 | 0.248 | 0.091 | 0.188 | 0.315 | 1.876 | 0.221 | 13.038 | 0.276 | 0.005 | 0.033 | 0.022 | 0.03 | 1.594 | 0.128 | 0.218 | 0.208 | 0.004 | 0.154 |
| **wt_20_C** | 0.14 | 0.099 | 0.036 | 0.099 | 0.157 | 0.704 | 0.088 | 2.579 | 0.108 | 0.004 | ND | 0.019 | 0.005 | 0.257 | 0.171 | 0.098 | 0.083 | 0.003 | 0.063 |
| **PKA1_100_A** | 1.759 | 0.234 | 4.149 | 0.554 | 1.312 | 1.881 | 0.308 | 23.783 | 0.445 | 0.014 | 1.007 | 0.279 | 10.522 | 6.332 | 0.425 | 0.397 | 0.271 | 0.067 | 0.51 |
| **PKA1_100_B** | 1.376 | 0.331 | 5.04 | 0.752 | 1.883 | 2.277 | 0.595 | 29.602 | 0.736 | 0.025 | 1.417 | 0.378 | 13.836 | 7.233 | 0.599 | 0.556 | 0.627 | 0.117 | 0.692 |
| **PKA1_100_C** | 1.443 | 0.274 | 4.636 | 0.77 | 1.875 | 2.174 | 0.533 | 27.585 | 0.652 | 0.02 | 1.339 | 0.355 | 13.052 | 7.049 | 0.581 | 0.548 | 0.521 | 0.11 | 0.697 |
| **PKA1_87_A** | 0.687 | 2.227 | 3.085 | 0.237 | 1.106 | 4.547 | 0.506 | 46.875 | 0.792 | 0.009 | 0.655 | 0.106 | 2.034 | 5.373 | 0.485 | 0.567 | 0.684 | 0.041 | 0.352 |
| **PKA1_87_B** | 0.532 | 1.088 | 3.855 | 0.214 | 1.315 | 3.529 | 0.399 | 26.884 | 0.681 | 0.021 | 0.575 | 0.132 | 2.243 | 6.127 | 0.476 | 0.453 | 0.414 | 0.039 | 0.34 |
| **PKA1_87_C** | 0.728 | 1.968 | 2.464 | 0.233 | 1.196 | 4.554 | 0.504 | 44.549 | 0.802 | 0.008 | 0.628 | 0.098 | 1.884 | 5.885 | 0.504 | 0.603 | 0.802 | 0.043 | 0.342 |
| **PKA1_50_A** | 0.474 | 4.13 | 2.421 | 0.202 | 0.279 | 3.624 | 0.467 | 53.792 | 0.813 | 0.009 | 0.263 | 0.057 | 0.531 | 4.992 | 0.333 | 0.449 | 0.733 | 0.036 | 0.261 |
| **PKA1_50_B** | 0.37 | 9.541 | 1.084 | 0.208 | 0.473 | 2.517 | 0.487 | 38.026 | 1.011 | 0.011 | 0.245 | 0.052 | 0.186 | 5.488 | 0.251 | 0.468 | 0.444 | 0.042 | 0.239 |
| **PKA1_50_C** | 0.518 | 4.934 | 2.292 | 0.223 | 0.362 | 4.586 | 0.669 | 52.862 | 0.858 | 0.008 | 0.313 | 0.054 | 0.468 | 4.465 | 0.355 | 0.48 | 0.868 | 0.043 | 0.296 |
